# Visual field map clusters in human frontoparietal cortex

**DOI:** 10.1101/083493

**Authors:** Wayne E. Mackey, Jonathan Winawer, Clayton E. Curtis

## Abstract

The visual neurosciences have made enormous progress in recent decades, in part because of the ability to drive visual areas by their sensory inputs, allowing researchers to reliably define visual areas across individuals and across species. Similar strategies for parceling higher-order cortex have proven elusive. Here, using a novel experimental task and nonlinear population receptive field modeling we map and characterize the topographic organization of several regions in human frontoparietal cortex. We discover maps of both polar angle and eccentricity that are organized into clusters, similar to visual cortex, where multiple maps of polar angle of the contralateral visual field share a confluent fovea. This is striking because neural activity in frontoparietal cortex is believed to reflect higher-order cognitive functions rather than external sensory processing. Perhaps the spatial topography in frontoparietal cortex parallels the retinotopic organization of sensory cortex to enable an efficient interface between perception and higher-order cognitive processes. Critically, these visual maps constitute well-defined anatomical units that future study of frontoparietal cortex can reliably target.

## INTRODUCTION

A fundamental organizing principle of sensory cortex is the topographic mapping of stimulus dimensions (Kaas, 1997; Mountcastle, 1957). For instance, visual areas contain maps of the visual field, whereby the spatial arrangement of an image is preserved such that nearby neurons represent adjacent points in the visual field (Holmes, 1918; Inouye, 1909). At a larger scale, multiple visual field maps are arranged in clusters, in which several adjacent maps share a common eccentricity representation (Kolster et al., 2009; Wandell et al., 2005; Wandell et al., 2007). These clusters are thought to form larger, more efficient processing units by sharing computational resources and minimizing the length of axons connecting the portions of the maps with similar spatial receptive fields (Wandell et al., 2005). Furthermore, it has been suggested that clusters, rather than individual visual field maps, organize specializations in cortical function (Bartels and Zeki, 2000). To date, more than twenty visual field maps have been identified in the human brain, several of which are organized into clusters (Arcaro and Kastner, 2015; Larsson and Heeger, 2006; Wandell et al., 2005; Wandell et al., 2007; Wandell and Winawer, 2011; Wang et al., 2015).

Recently, using modified versions of the standard traveling wave method for mapping early visual cortex, several labs have identified maps of polar angle along the intraparietal sulcus (IPS) in posterior parietal cortex and the precentral sulcus (PCS) in frontal cortex (Jerde et al., 2012; Kastner et al., 2007; Schluppeck et al., 2005; Sereno et al., 2001; Silver et al., 2005; Swisher et al., 2007). Not surprisingly, these maps have attracted a great deal of attention given their presumed involvement in a wide range of cognitive and sensorimotor processes, including attention, working memory, and decision-making (Bechara et al., 1994; Mackey et al., 2016a; Mackey et al., 2016b; Manes et al., 2002; Posner et al., 1984; Wilkins et al., 1987). Some of these visual maps may correspond to the human homologs of well-characterized areas in the macaque brain that are topographically organized, like the lateral intraparietal area (LIP) and the frontal eye field (FEF). Yet, such inter-species homology and how these maps contribute to different aspects of behavior and cognition remain unknown. This is likely due to our limited understanding of the basic organizing principles of these maps. Further challenges are posed because the maps in parietal and especially frontal cortex have less reliable stimulus-evoked BOLD signals compared to visual cortex, are more coarsely organized, and show less consistent topography across subjects. In some cases it is uncertain as to whether a region merely has a contralateral bias as opposed to containing an actual topographic map (Hagler and Sereno, 2006; Jerde et al., 2012; Kastner et al., 2007; Patel et al., 2014; Silver and Kastner, 2009). Here, we take a step back from trying to understand the functions and instead systematically characterize the basic organizing principles of these putative visual field maps in frontoparietal cortex.

Establishing the organization of maps in parietal and frontal cortex will have several major impacts. First, identification and better characterization of which maps are human homologs of macaque areas will facilitate better translation of non-human primate models of human cognition to humans. Second, understanding the topography of parietal and frontal maps will enable researchers to aggregate results at the level of individual maps or even small areas within maps, rather than at the level of large regions of cortex, similar to the successes in delineating maps in occipital cortex in animals (Desimone and Ungerleider, 1986; Gattass et al., 2005; Van Essen and Zeki, 1978) and humans (Sereno et al, 1995; Engel et al 1997). Indeed, the fundamental reason the *visual* neurosciences have outpaced the *cognitive* neurosciences is the ability to reliably define and study the function of the same areas across individuals and across labs. Such convention facilitates comparisons between studies of different computations and representations across different subject populations and methods of measurement. This effectively creates a worldwide, across-time, collaboration between all labs. As such, the organization of visual field maps in early visual cortex has been well characterized, to the point of enabling the development of detailed templates of the visual field for V1-V3 (Benson et al., 2014; Benson et al., 2012; Dougherty et al., 2003). In contrast, much less is known about the organization of visual field maps in frontoparietal cortex, and thus presents a critical roadblock for understanding their functions.

To these ends, we focus on characterizing the organization and retinotopic properties of putative visual field maps in frontoparietal cortex. We estimated population receptive field (pRF) parameters in topographic areas in early visual, parietal, and frontal cortices. The pRF method not only estimates a voxel’s polar angle and eccentricity preference, but also its receptive field (RF) size, and has been shown to more accurately map topography than conventional traveling wave methods (Dumoulin and Wandell, 2008). However, two important challenges exist when attempting to identify visual field maps in frontoparietal cortex. First, passive viewing of high-contrast spatial patterns elicits weak and non-systematic responses in frontoparietal cortex (Saygin and Sereno, 2008; Silver et al., 2005). Therefore, identifying visual field maps in higher order frontoparietal cortex requires more cognitively demanding stimulation that taxes attention or memory (Jerde et al., 2012; Silver et al., 2005). Second, the large RF sizes expected in frontoparietal cortex make it difficult for linear RF models to accurately characterize response properties to stimuli that vary in size. To overcome these challenges, we developed a novel, attention-demanding task specifically designed to elicit robust, systematic responses in frontoparietal cortex (Figure 1A). Moreover, we estimated pRFs with a model that accounts for nonlinear responses to stimuli of varying size (Kay et al., 2013; Winawer et al., 2013). Nonlinear spatial summation is more pronounced in extrastriate maps than in V1 and is likely to be even more so in frontoparietal cortices.

## RESULTS

During a single functional neuroimaging session, observers swept their focus of attention across the visual field, while maintaining central fixation, in order to perform a difficult motion discrimination task. The task was designed to tax attentional resources presumably controlled by activity in topographic maps in frontoparietal cortex (Figure 1A). A bar aperture swept across the visual field in discrete steps, vertically or horizontally traveling in four possible directions: left to right, right to left, top to bottom, bottom to top. The bar was comprised of three rectangular patches, each of which contained a random dot kinematogram (RDK) moving in a particular direction. The central patch contained 100% coherent motion and the two flanking patches contained low coherence motion. Observers pressed a button to indicate which of the two flanking patches (above or below for vertical bars, left or right for horizontal bars) matched the RDK direction of the middle patch. The motion coherence of the flanking patches was staircased to ensure the task remained difficult throughout the duration of the scanning session (75% accuracy).

**Figure 1:**
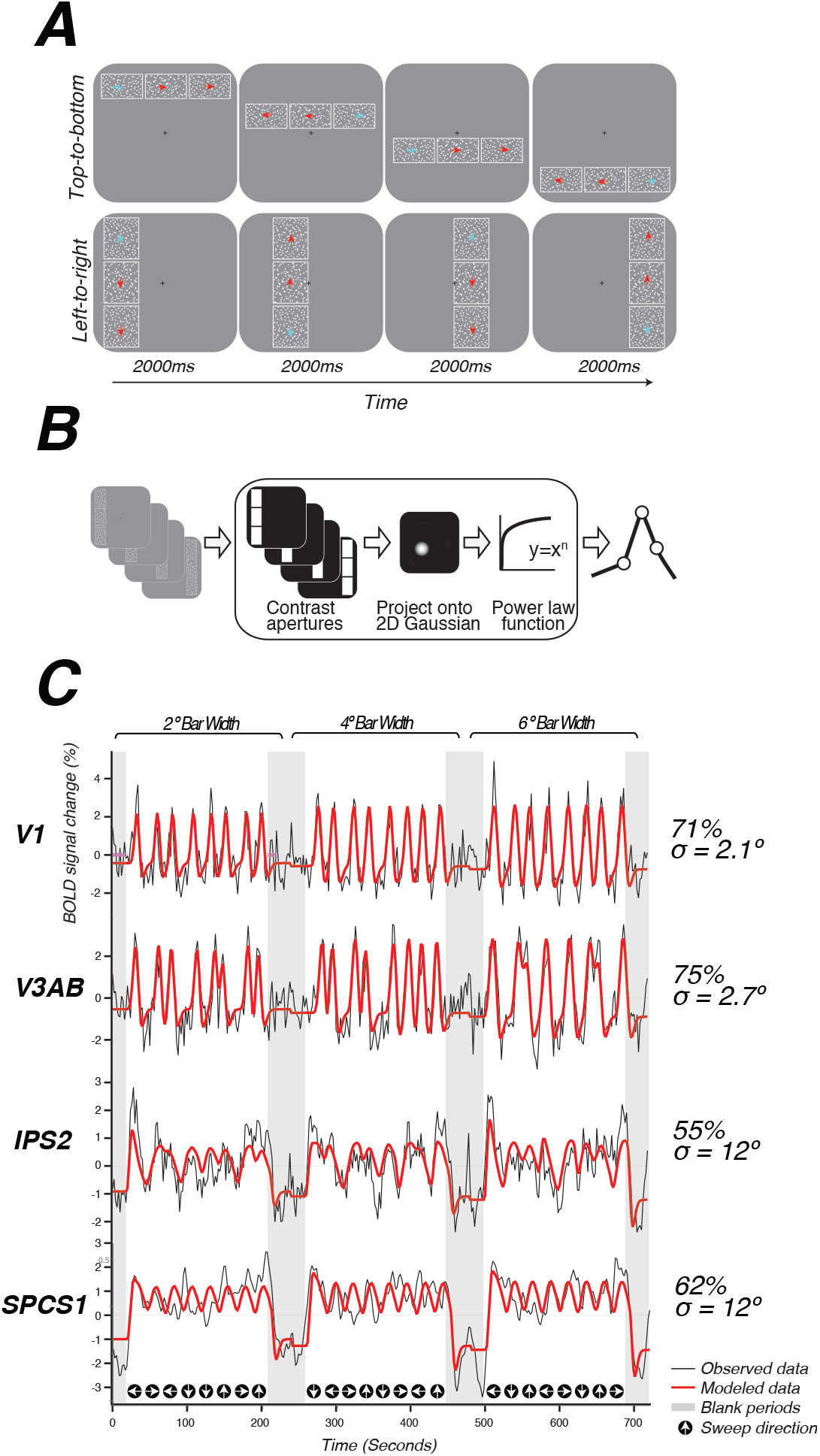
Topographic mapping and population receptive-field modeling. (***A***) Discrimination task used for topographic mapping. Subjects fixated at the center of the screen while attending covertly to a bar composed of three apertures of moving dots incrementally traversed the screen. Subjects indicated on each trial which aperture (left, right, top, or bottom) was comprised of dots whose motion direction matched the motion direction of the dots in the middle sample aperture. Motion coherence was staircased in order to constantly tax attention. The white outlines around each of the three apertures are shown here for clarity, but were not visible to subjects. (***B***) Schematic of the nonlinear population receptive-field modeling procedure. Trial sequences were converted into 2D binary contrast apertures and projected onto a 2D Gaussian representing a predicted pRF. Static non-linearity was applied to account for compressive spatial summation. (***C***) Example model fits across multiple visual field maps. pRF model *Observed data Modeled data Blank periods Sweep direction* predictions are shown in red, actual data for an individual voxel for a given visual field map is shown in black. Stimulus sweep direction and bar width are shown above and below the model fits. Estimated pRF size and variance explained for each voxel is shown to the right.

We used a nonlinear pRF model to predict the BOLD response of each individual voxel to the visual stimulus (Figure 1B). Each model is specified by the center (x,y) and size (sigma, or standard deviation of isotropic 2D Gaussian) of the pRF, and a power-law exponent (n) to account for sub-additive spatial summation (Equations 1&2). The non-linearity interacts with the Gaussian standard deviation to make an effective pRF size of 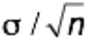 Kay et al., 2013; Winawer et al., 2013). We excluded voxels from further analysis if less than 10% of the variance in the time series was explained by the pRF model (Equation 3). We also excluded voxels with pRF centers outside of the limits of our visual display (12 degrees of visual angle).

The pRF fitting strategy proposed by Dumoulin and Wandell (2008) was a coarse-to-fine approach, in which the initial coarse fit was solved on time series that were spatially blurred (approximating a Gaussian kernel of 5 mm width at half height) and temporally decimated (2x), and used gridded parameters rather than searching for the best fit using nonlinear search optimization algorithms; the second stage then used the solution of the grid fit as a seed for a search, and applied this search to the unblurred (in space and time) time series. Here, we used only the first stage (the grid fit), applied to the smoothed and temporally decimated time series. The grid fit is more robust to noise (though also less accurate when noise is low). Because our primary goal was to map frontoparietal cortex, where pRFs were expected to be larger, and visually driven signals were expected to be less reliable than visual cortex, the grid fit was chosen. The grid included the same set of possible values in the solution as described in Dumoulin and Wandell, with the addition of the power law exponent, which was gridded to be 0.25, 0.5, 0.75, or 1. A value of 1 indicates a linear fit; values less than 1 indicate increasingly more sub-additive spatial summation.

We identified retinotopic maps in early dorsal visual areas (V1, V2, V3, V3AB) as well as at least four maps along the intraparietal sulcus (IPS0, IPS1, IPS2, IPS3) and two regions with spatial tuning along the precentral sulcus in frontal cortex in each hemisphere of all subjects. The superior portion of the precentral sulcus region contained two distinct visual field maps that we refer to as SPCS1 and SPCS2. Example model fits for voxels in different visual field maps are shown in Figure 1C. Although we were able to observe additional parietal maps in some subjects, we restricted further analysis to maps consistently observed in both hemispheres of every subject. All areas are described in further detail below.

### Visual cortex

As an important control, our methods revealed the functional organization and pRF properties typically observed in early visual cortex. Polar angle and eccentricity maps revealed the expected patterns in V1, V2, V3, and V3A&B (Figure 2A&B). The progression of these areas begins medially from the occipital pole in the calcarine sulcus with V1, and extends dorsally and laterally along the surface of the cortex. V1 contains a continuous angular representation of the contralateral hemifield, beginning at the upper vertical meridian (UVM) on the lower bank of the calcarine sulcus, and progressing to the lower vertical meridian (LVM) along the inferior-to-superior direction. V2 sits adjacent to the LVM border of V1, and begins a reversal of phase angles representing half of the contralateral hemifield and progressing to the horizontal meridian (HM). V3 sits adjacent to the HM border of V2, beginning another phase reversal back towards the LVM, and also represents half of the contralateral hemifield. Finally, V3A begins along the border of V3 in the periphery (but not fovea), and contains a full and continuous angular representation of the contralateral hemifield from the LVM to the UVM. V3B is adjacent to V3A, divided by a shared foveal representation, as reported previously (Press et al., 2001; Wandell et al., 2005).

Together, these maps form two distinct visual field map clusters. A cluster is comprised of a group of angle maps that all share a confluent fovea (Kolster et al., 2009; Wandell et al., 2005). Within a cluster, the boundaries of adjacent angle maps can be defined by reversals in polar angle progression or eccentricity. For example, the boundary between V1 and V2 is identified by a polar angle reversal at the LVM, as described above. V1, V2, and V3 share a common foveal representation centered near the occipital pole that extends towards the collateral sulcus. V3A and V3B comprise a second cluster because they share another foveal representation, anterior to and distinct from the foveal representation shared by V1, V2, and V3. They are divided by an eccentricity reversal rather than a polar angle reversal.

Moving up the hierarchy of early visual cortex, from V1 to V3AB, pRF parameters begin to differ systematically. For example, pRF sizes at a given eccentricity generally increased across visual field maps, and within each visual field map in visual cortex, pRF size increased monotonically with eccentricity (Figure 2C). Consistent with previous studies investigating spatial summation in pRFs (Kay et al., 2013), we found that in extrastriate maps, the pRF models show more sub-additive spatial summation, indicated by a smaller power law exponent. The fraction of voxels with the minimal pRF exponent allowed by our grid search (0.25) was least in V1 (33%), and substantially higher in extrastriate areas (68% in V2; 80% in V3; 77% in V3A&B)

**Figure 2:**
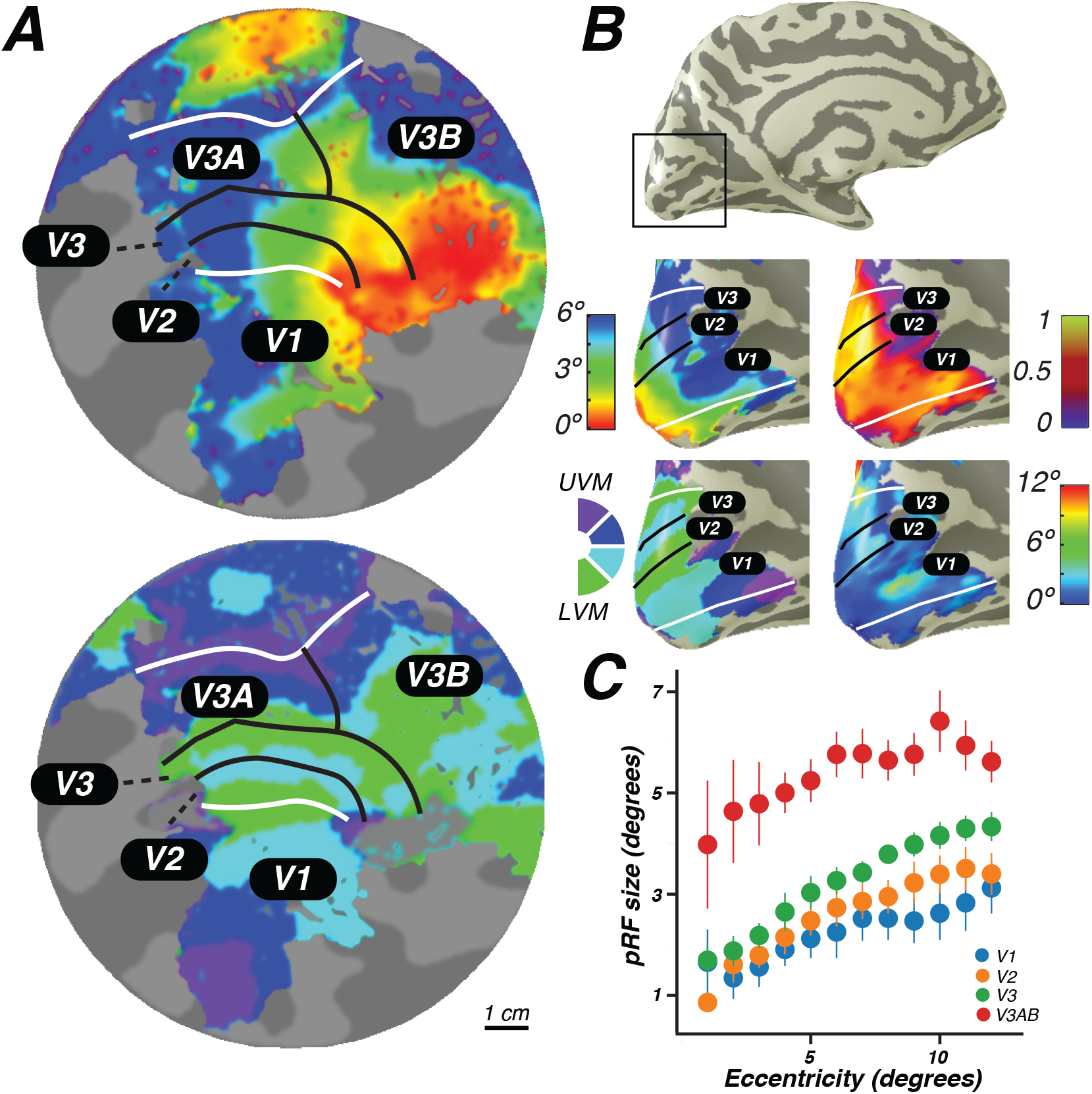
Visual field maps in early visual cortex. The color of each voxel indicates the best fit pRF parameter for the data being displayed. (***A***) Eccentricity (top) and polar angle (bottom) maps in the left hemisphere of an example subject projected on a flattened representation of the cortical surface, where dark gray denotes sulci and light gray denotes gyri. V1, V2, and V3 share a confluent fovea while V3A and V3B share another. (***B***) Eccentricity (top left) and polar angle (bottom left) maps, along with variance explained (top right), and pRF size (bottom right) of an example subject projected on an inflated cortical surface. (***C***) Relationship between pRF size and eccentricity. pRF sizes of voxels in V1, V2, V3, and V3AB increase with eccentricity. Error bars represent ± 1 SEM across subjects. Individual subject maps are shown in **Figure S1.**

### Parietal cortex

We found clear patterns of systematic organization in eccentricity and polar angle maps in the intraparietal sulcus (Figure 3). Beginning with IPS0 and progressing through IPS3, each map contains a full representation of the contralateral hemifield, while the polar angle reversals demarcate the boundaries of each individual map. Additionally, each successive map lies anterior to the visual field map before it. Starting with the most posterior map, IPS0 lies at the intersection of the transverse occipital sulcus and the intraparietal sulcus, adjacent to the UVM representation of V3AB. The angular representation systematically progresses from the UVM to the LVM, where another phase reversal takes place, creating the posterior border of IPS1. The angular map of IPS1 sweeps from the LVM back towards the UVM, where it borders IPS2. IPS2 then contains an angular progression from the UVM to the LVM, where it borders IPS3. IPS3 then contains an angular progression from the LVM to the UVM.

Using these novel methods, we discovered that maps along the IPS are organized into two clusters of visual maps. IPS0 and IPS1 share a confluent fovea, while IPS2 and IPS3 share another, distinct foveal representation (Figure 3, **second column**). However, unlike maps in early visual cortex, we were unable to measure pRF sizes in parietal visual field maps with sufficient accuracy. Estimates of pRF size for nearly all voxels in parietal visual field maps were ~12 degrees, even for voxels with pRF centers at the fovea. As 12 degrees is both the maximum stimulus extent in our experiment and the upper boundary of our grid fit, it is likely that these pRF size estimates are a floor on the true size, rather than an accurate measure of the pRF size of voxels in this region. However, these measurements do indicate that the sizes of pRFs in parietal cortex are large, and likely larger than 12 degrees.

**Figure 3:**
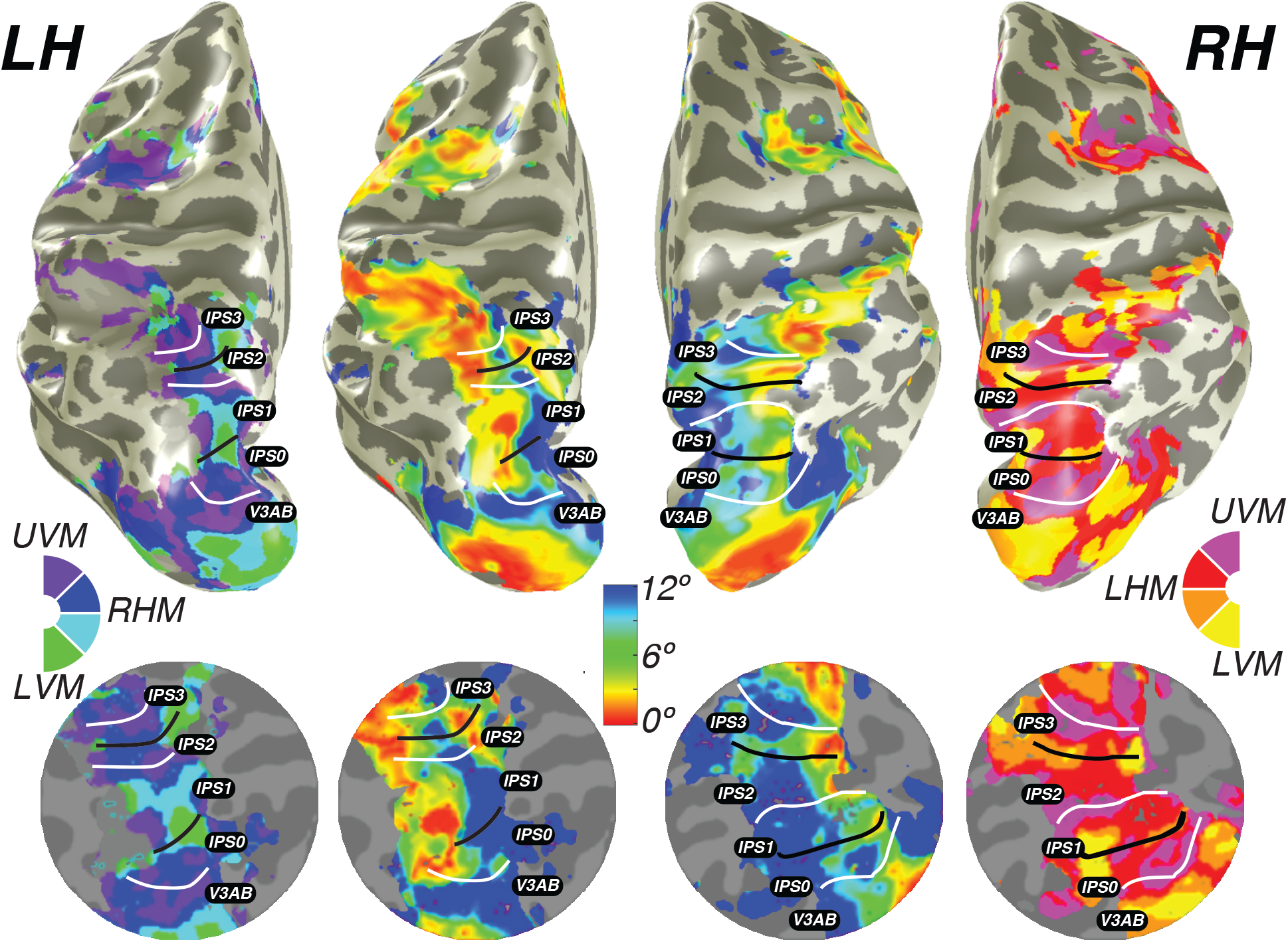
Visual field maps in parietal cortex. Maps of polar angle and eccentricity on an inflated cortical surface (top and bottom) and on a flattened representation of the cortical surface (middle) for an example subject. IPS0/IPS1 form one visual field map cluster, while IPS2/IPS3 form another. Each cluster consists of two angle maps that share a confluent foveal representation. White lines denote the boundaries at the upper vertical meridian (UVM) and black lines denote the lower vertical meridian (LVM). Individual maps for all subjects are provided in supplementary materials (**Figure S2**).

Maps in parietal cortex, like extrastriate maps in visual areas, showed a systematic sub-additivity in spatial summation. The vast majority of voxels were best fit by a pRF model with the minimal pRF exponent allowed by our grid (IPS0: 73%; IPS1: 75%; IPS2: 81%; IPS3: 89%).These results are comparable to the low exponent found in measurements of ventral occipitotemporal face-selective regions (0.20, 0.16, 0.23 in three face-selective ROIs)(Kay et al., 2015).

### Frontal cortex

We also discovered that visual field maps in frontal cortex are organized by polar angle as well as eccentricity. Although polar angle maps have been described before (Jerde et al., 2012; Kastner et al., 2007; Saygin and Sereno, 2008), maps of eccentricity have remained undiscovered until now. Two regions of frontal cortex along the PCS exhibited spatial tuning: one in the superior portion (SPCS) and another in the inferior portion (IPCS) (Figure 5). Each region contains a map of eccentricity with a central foveal representation that radiates towards the periphery. In the SPCS, the fovea is represented in the fundus of PCS where it intersects the superior frontal sulcus. The fovea representation in the IPCS is located at the junction of the PCS and the inferior frontal sulcus. This organization was consistent across both hemispheres in all subjects (Figure 4). The discovery of the foveal representation in a location common across subjects allowed us to separate the region into two visual field maps, identifiable in each subject. We refer to these new maps as SPCS1 and SPCS2 consistent with naming visual field maps according to their anatomical location (Arcaro et al., 2009; Larsson and Heeger, 2006). Figure 5 shows maps of angle and eccentricity projected on both flattened and inflated cortical reconstructions. Both SPCS1 and SPCS2 contain a full and continuous representation of the contralateral hemifield. The superior border of SPCS1 begins at the LVM and continues to the UVM, where a phase reversal occurs signaling the border of SPCS2. SPCS2 then contains an angular progression from the UVM back to the LVM.

**Figure 4:**
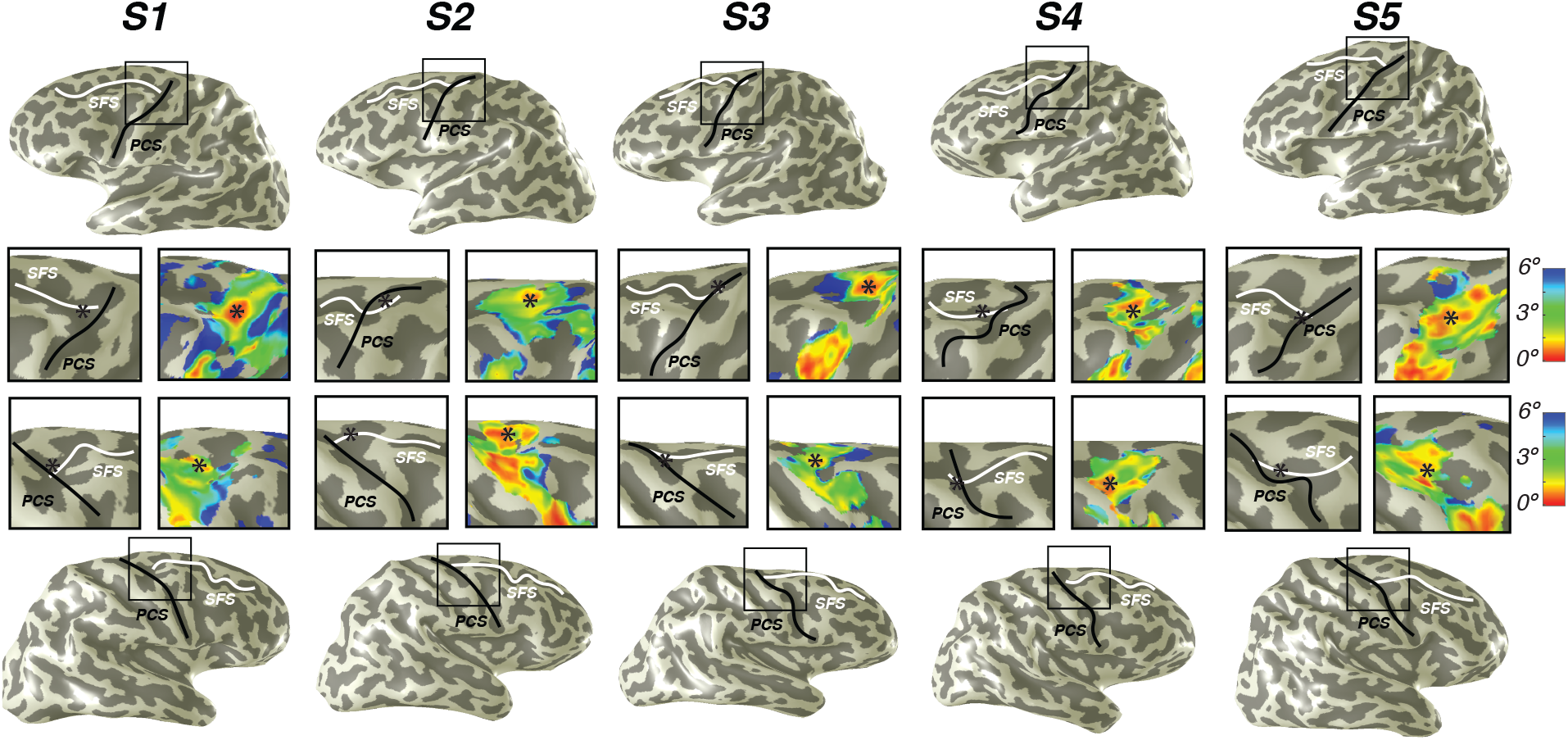
Representation of fovea in SPCS map. Left (top) and right (bottom) hemispheres for each individual subject are shown. Top and bottom rows mark the anatomical locations of the superior frontal sulcus (SFS: white line) and superior precentral sulcus (SPCS: black line) on an inflated cortical surface representation of each hemisphere. For clarity, black squares represent a zoomed in view of the anatomical intersection of the SFS and SPCS. The location of the fovea (asterisk) is shown both on the raw anatomy (left), as well as the eccentricity map (right) for each individual subject. Notice how the fovea lies at the intersection of the SFS and SPCS for each subject.

The inferior portion of the PCS (IPCS) also contained spatial representations, and was distinct from the SPCS cluster. Although the topography in the IPCS region was not sufficiently regular across subjects, peripheral representations typically surrounded a foveal representation. Although less clear, the visual field coverage showed a systematic representation of the contralateral visual field, similar to SPCS1 and SPCS2, which we return to in the next section (*Visual field coverage density and laterality*).

As was the case in parietal cortex, in frontal cortex, estimates of pRF size were large and pRF exponents small for nearly all voxels. Size estimates were limited by the field of view our display and the analysis method. Most pRFs in frontal cortex had a size estimate at the upper bound of our analysis (12 deg) and an exponent at the lower bound (0.25; IPCS: 78%; SPCS1: 82%; SPCS2: 75%) As discussed below in the next two sections, the large pRF size does not indicate a failure to estimate spatial tuning, as indicated by the cross-validated model fits and the laterality indices.

**Figure 5:**
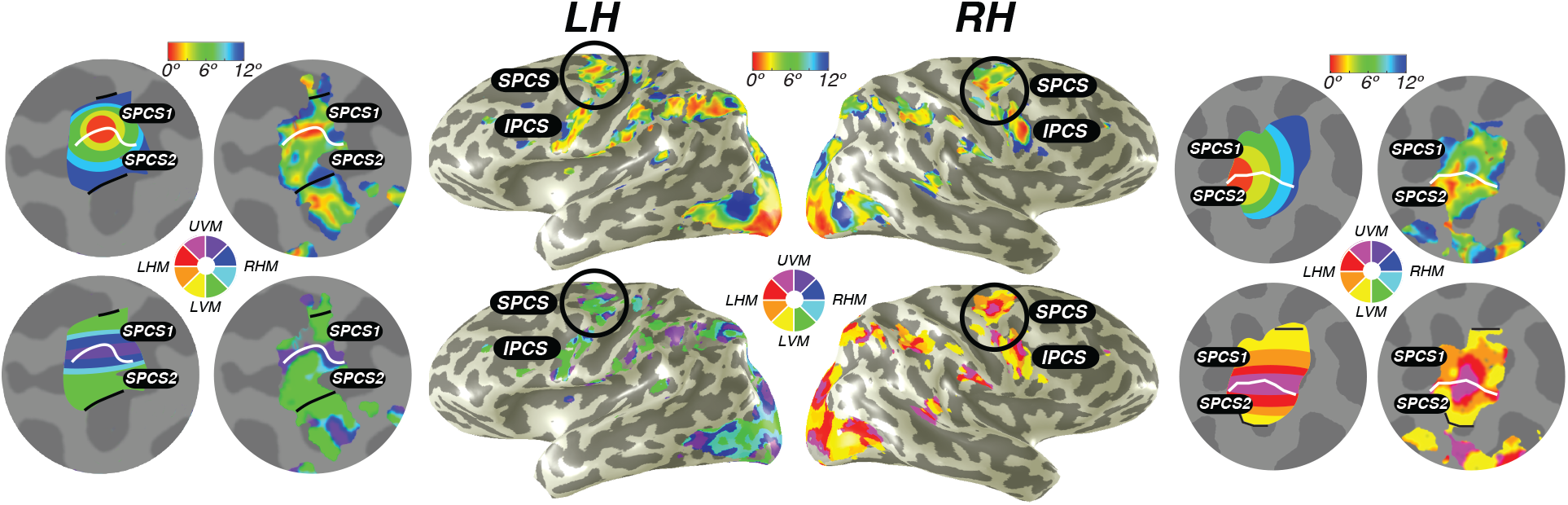
Visual field maps in frontal cortex. Maps of eccentricity and polar angle are displayed for an example subject projected on both inflated (inside) and flattened (outside) cortical surfaces. In order to clearly demonstrate the systematic organization of each map, each pair of flattened cortical surfaces depicts a cartoon schematic of the organization of each map (left column) next to actual map data (right column). SPCS1 and SPCS2 form a visual field map cluster, sharing a foveal representation that sits at the intersection of the PCS and the SFS. White lines denote the boundaries at the upper vertical meridian (UVM) and black lines denote the lower vertical meridian (LVM). The foveal representation in IPCS sits at the intersection of the PCS and the IFS. Individual maps for all subjects are provided in supplementary materials (**Figure S3**).

### Visual field coverage laterality and density

A pRF model summarizes the sensitivity of a single cortical site (e.g., a voxel) to positions in the visual field. By combining the pRFs across sites within a region of interest, one can visualize the field of view of the region of interest, also called the visual field coverage (Amano et al., 2009; Wandell and Winawer, 2015; Winawer et al., 2010). The visual field coverage is typically computed as the envelope of the pRFs within an ROI. Here, we took the mean of the pRFs rather than the envelope, thereby scaling the visual field coverage by the density of pRFs at any particular location in the visual field (cortical magnification), which we refer to as the coverage density. In the V1-V3 maps, the coverage density plots show a nearly total contralateral bias, as well as bias toward the fovea over the periphery (Figure 6A). The contralateral and foveal biases in early visual cortex reflect three aspects of the pRF models: (1) the centers are in the contralateral visual field, (2) the pRF sizes are relatively small, and (3) the largest number of pRF centers are close to the fovea. In V3AB, the coverage density is wider and extends slightly into the ipsilateral visual field, resulting from larger pRF sizes. This observation was used previously to distinguish area MST from area MT in the human visual system (Amano et al., 2009; Huk et al., 2002).

In parietal cortex (IPS0-IPS3) and frontal cortex (IPCS and SPCS1&2), the coverage is also centered in the contralateral hemifield, but increasingly extends into the ipsilateral field, again reflecting the larger pRF sizes. Similar to visual cortex, the coverage density is highest near the fovea, indicating a qualitative similarity in cortical magnification between frontoparietal cortex and visual cortex.

We quantified the degree to which visual areas have lateralized pRFs using a laterality index (Equation 4). In agreement with the coverage density plots, the laterality index shows that early visual cortex is highly lateralized, while successive maps from parietal to frontal cortex become less and less lateralized (Figure 6C). Additionally, there is larger subject-to-subject variability in the lateralization index in parietal and frontal areas compared to visual cortex. In sum, although maps in frontal and parietal cortex have very large pRFs compared to visual cortex, the spatial tuning is sufficiently reliable to show clear lateralization, complete hemifield visual coverage, and maps that are organized in an orderly topographic manner.

**Figure 6:**
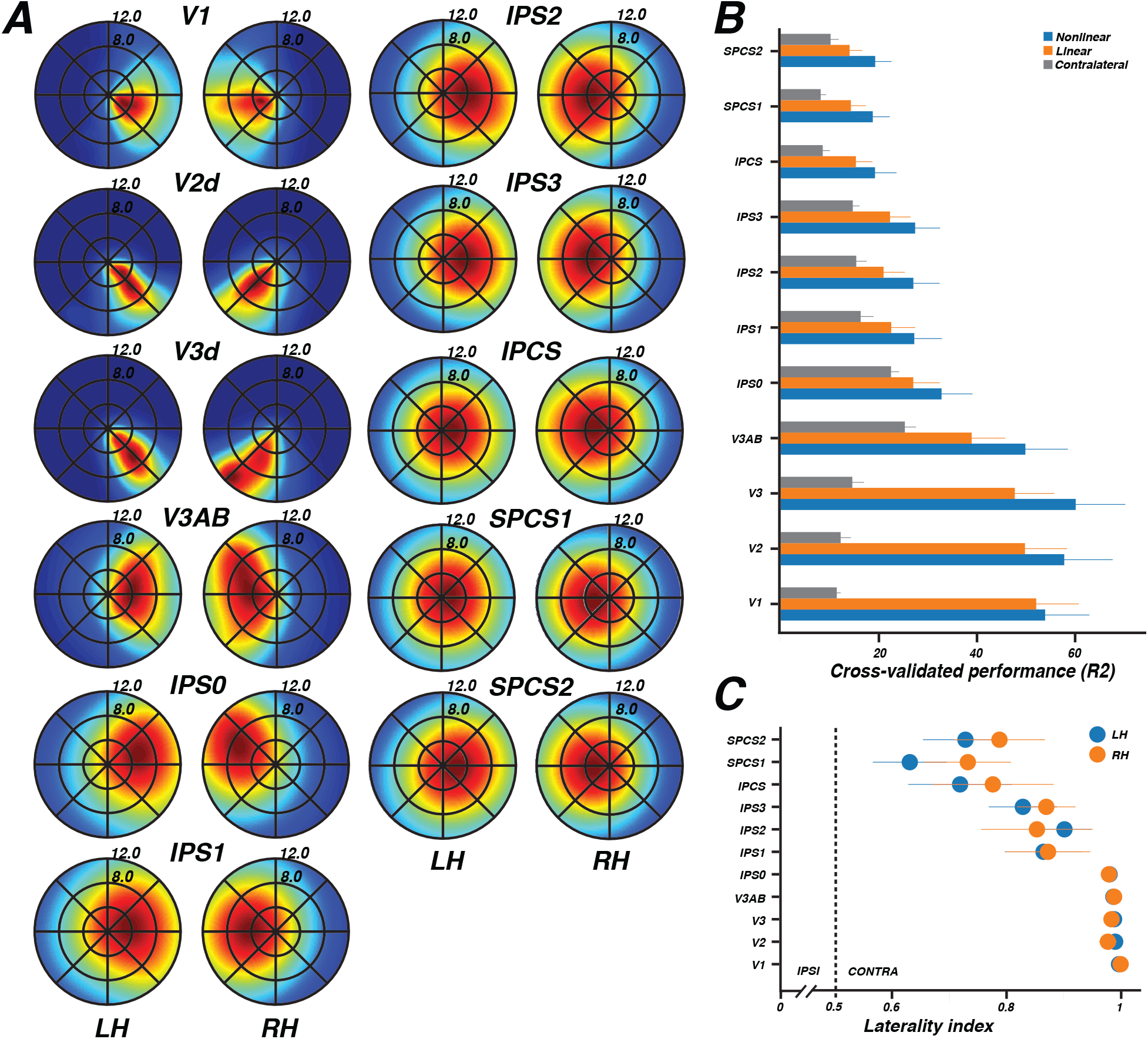
Visual field map coverage, laterality, and model comparison. (***A***) Visual field map coverage plots. While all visual field maps primarily represent the contralateral hemifield, maps in association cortices begin to also represent small portions of the ipsilateral hemifield perifoveally. This is due to the fact that pRFs are less eccentric and larger in frontal and parietal cortex than early visual cortex. (***B***) Comparison of cross validation results by model (Means ± 1 SEM across subjects). For every visual field map, the non-linear model explained the largest amount of variance, followed by the linear model, and finally the contralateral model. (***C***) Laterality index (Means ± 1 SEM across subjects). The index ranges from 0 (completely ipsilateral) to 0.5 (no laterality) to 1 (completely contralateral). All areas are highly contralateralized. Early visual cortex shows very little variability across subjects, while frontal and parietal cortices exhibit some variability across subjects.

### Model comparison

Reliably mapping topography in frontal and parietal cortex depends on nonlinear models of pRFs. We compared performance of the nonlinear pRF model to two other models: a linear pRF model (Dumoulin and Wandell, 2008) and a simple contralateralized response model. Comparing with the linear model allowed us to see how much, if any, improvement in accuracy was gained by allowing for sub-additive spatial summation. Comparing with the lateralized response model allowed us to investigate whether our results indicate systematically organized maps as opposed to noisy representations of lateralized responses, with no spatial tuning other than a preference for the contralateral hemifield. The lateralized model predicted a uniform response amplitude whenever any portion of the stimulus was in the contralateral visual field, and zero response otherwise. In order to compare model performance, we used a leave-one-out cross-validation procedure, which provides no advantage to models with additional parameters. The models were solved using two of the three stimulus types as training data (narrow, intermediate, or wide bars), and the remaining stimulus type as test data, iterated by leaving out each of the three stimulus types. We defined accuracy as the variance explained for the left out data, averaged across the three cross-validation iterations. The ordinal ranking of the three models was the same in all 11 areas tested: the nonlinear model explained the greatest amount of variance, followed by the linear model, and then the lateralized response model (Figure 6B). The quantitative advantage of the nonlinear model over the linear model was smallest in V1, and larger in other maps. The improvement in performance from the nonlinear over the linear model demonstrates the benefits of including a parameter to estimate sub-additive spatial summation. This is particularly true for maps in frontal and parietal cortex, where pRFs are large and response nonlinearity estimations were at our measurement boundary. The contralateral model was much worse than either of the Gaussian models (linear or nonlinear) in visual cortex, a result of the relatively small pRFs in those areas, yet even in parietal and frontal areas, where pRFs were much larger, the contralateral model was worse than the non-linear pRF model in every area tested. This result strengthens the claim that the observed responses are spatially tuned, and not merely due to a non-specific preference for contralateral stimuli. These observations, coupled with our newly discovered systematic organization of frontal and parietal maps, demonstrate the existence of topographically organized visual field maps in frontoparietal cortex.

## DISCUSSION

Using novel procedures, we precisely characterized the topographic organization of visual field maps in human frontoparietal cortex, including four visual field maps along the IPS and two spatially-tuned regions along the PCS. Each of these maps contains a representation of the full range of polar angles in the contralateral visual field; they are topographically, not simply contralaterally, organized. Combining pRFs across voxels within visual maps, we demonstrate that these maps tile the complete contralateral hemifield in an orderly manner. As expected, the pRF sizes in frontoparietal cortex are larger than those in early visual cortex. We also demonstrate clear maps of eccentricity along all IPS, which to date has only been reported in a single subject (Swisher et al., 2007), and in PCS maps, which have never been reported. Furthermore, we show that a spatially tuned region of the superior precentral sulcus is organized into at least two distinct visual field maps (referred to here as SPCS1 and SPCS2), each representing the entire contralateral visual field in an orderly manner. Interestingly, the visual maps in both parietal and frontal cortex are organized into clusters of polar angle maps sharing the dimension of eccentricity, similar to visual cortex (Wandell et al., 2005). Two maps in frontal cortex and two pairs of maps in parietal cortex form clusters of polar angle maps that share a foveal to peripheral representation. Together, these data clearly describe the topographic structure of the visual maps in human frontoparietal cortices.

### Frontoparietal cortex, like visual cortex, is organized into clusters

We demonstrate that visual field maps in parietal and frontal cortex are organized into map clusters, similar to the organization found in early visual cortex. Visual cortex is composed of four to five distinct visual field map clusters (Wandell et al., 2007). Individual maps in these clusters may perform similar, yet distinct computations that together form a larger processing unit. Their close proximity and short-range connections make information processing more efficient. We propose the existence of at least 3 clusters along human IPS and PCS. Following the pattern starting with V3AB, we find two pairs of angle maps sharing a confluent foveal to peripheral representation: IPS0/IPS1, and IPS2/IPS3.

Based on the report from a single subject, Swisher et al. (2007) hinted that a foveal representation might be shared by IPS0 and IPS1 positioned on the medial wall of the IPS that progressed laterally towards the periphery. We find such an organization to be inconsistent across subjects. Although some subjects had this mediolateral organization, others show the opposite – a foveal representation on the lateral wall of the IPS (e.g., S5 in Figure 3) or in the fundus of the IPS. Along the PCS, we find a map cluster in the superior portion (SPCS1/SPCS2) sharing a common foveal representation at the border of the two maps.

While the non-human primate brain is typically investigated as a model of the human brain, it is important to note that the stimulus selectivity and the number and organization of visual field maps are not identical in human and non-human primates (Tootell et al., 1997; Wandell and Winawer, 2011; Winawer et al., 2010), and likely differ even more substantially in frontoparietal cortex. For example, although human IPS contains the putative homologue of the monkey lateral intraparietal area (LIP), human IPS houses at least four individual visual field maps (IPS0, IPS1, IPS2, IPS3) organized into two clusters (IPS0/IPS1 and IPS2/IPS3). Macaque LIP however appears to only contain a single visual field map, with no consistent organization of eccentricity (Arcaro et al., 2011; Ben Hamed et al., 2001; Blatt et al., 1990)(but see (Patel et al., 2014). While it is likely that some of these inconsistencies between the species are the result of evolution and the superior cognitive capabilities of humans (Passingham, 2009; Rilling, 2006), it is also possible that additional macaque visual maps remain undiscovered. Although labs have mapped visual areas in non-human primates using traditional phase-based retinotopic methods (Arcaro et al., 2011; Kolster et al., 2009; Patel et al., 2010; Savaki et al., 2010), it is entirely possible that computational neuroimaging approaches, like those used here, might reveal additional visual field maps in non-human primates and in turn help bridge translational research.

How and why the brain contains so many redundant maps of visual space are longstanding puzzles (Barlow, 1986). Certain maps, such as V1 and MT, may serve to anchor the development of other visual field maps towards clusters (Rosa, 2002). Perhaps the map cluster in early visual cortex acts as a developmental blueprint for replication in frontoparietal cortex (Buckner and Krienen, 2013; Rosa, 2002). Indeed, the topographic development of visual areas without retinal input mimics the organization of earlier visual areas (Bourne and Rosa, 2006; Rosa and Tweedale, 2005). Therefore, similar to V1 and MT in sensory cortex, we suggest that visual field map clusters in IPS and PCS may serve as organizational anchors in the development of frontoparietal cortex.

### New visual field maps within the PCS form a visual field map cluster

We report a new subdivision of the human superior PCS consisting of two distinct angle maps (SPCS1 and SPCS2) that share a confluent foveal to peripheral representation, thereby forming a visual field map cluster. Each map contains a full representation of the contralateral visual field. A reversal in the phase angle demarcates the border between SPCS1 (LVM to UVM) and SPCS2 (UVM to LVM). The foveal representation shared by both maps sits at the intersection of the SPCS and the superior frontal sulcus. Together, these observations indicate that the eccentricity map and the angle map are not perfectly orthogonal, differing from V1, V2, and V3. However, nor are the two maps perfectly parallel, allowing the area to represent the entire contralateral hemifield. This is similar to the observation in LO1 and LO2, which also deviate from orthogonality between the angle map and eccentricity map, but nonetheless appear to each represent a full hemifield (Larsson and Heeger, 2006). The organization of the SPCS into two maps, while never before reported in humans, aligns closely with recent topographic mapping results in non-human primates (Savaki et al., 2015) and studies of functional connectivity and domain specialization in humans (Michalka et al., 2015; Power et al., 2012; Wig et al., 2014).

The SPCS is the putative homologue of the monkey frontal eye fields (FEF) (Blanke et al., 1999), which reside in the arcuate sulcus of the macaque (Bruce et al., 1985). Low-level electrical stimulation of macaque FEF neurons reliably elicits saccades to particular locations in the visual field (Bruce et al., 1985; Robinson and Fuchs, 1969). Despite this well-known property of FEF neurons, the topographic organization of macaque FEF is poorly understood. For instance, studies have revealed a coarse map of saccade amplitude where small saccades are represented in the ventral portion of the FEF and increasingly larger saccades are represented progressively more dorsal. However, the reported correspondence between stimulation site and saccade direction do not form a clearly organized map of the visual field (Bruce et al., 1985; Robinson and Fuchs, 1969). Two possibilities exist for the observed discrepancy between the organization of the macaque FEF and the organization of the SPCS we describe here. First, it may be that the human homologue of the macaque FEF has expanded to contain multiple maps and a more systematic topographic organization. Such differences should be expected given the 25 million years of evolutionary divergence between monkeys and humans (Blair Hedges and Kumar, 2003). Second, the coarse spatial resolution of neuroimaging may be more suited than precise electrical stimulation when quantifying the large-scale topographic organization. A recent study suggests that this may be the case. Savaki and colleagues (2015) imaged the distribution of metabolic activity in the macaque FEF and discovered two topographic maps of saccades to the contralateral visual field. Similar to what we observed in our data, the maps were separated by the vertical representation of space and each contained a visuomotor map of the entire contralateral visual field. The dorsal part of the arcuate sulcus contained a representation of the LVM, and progressed ventrally to a representation of the UVM. While the visuomotor maps in the monkey arcuate sulcus progressed along the rostral-caudal axis, the maps we observe progress along the dorsal-ventral axis. It is possible that this is simply a difference between the species, as other cortical (Orban et al., 2004) and subcortical (Arcaro et al., 2015) visual field map clusters are organized differently between the two species.

Our results help shed new light on a claim that maps in frontal cortex are organized differently than maps in parietal or visual cortex (Silver and Kastner, 2009). The primary evidence supporting this conclusion was that the same point in space was represented multiple times in the SPCS, while a given point in space was represented only once in a given map in parietal or visual cortex. We argue that this apparent discrepancy stems from treating the SPCS as a single map, rather than as two maps (SPCS1 and SPCS2), as we propose here. Indeed, if one were to combine any two smaller maps in parietal or visual cortex, the same point in space would be represented more than once. Although not discussed in the original papers, a few subjects appear to have a phase reversal in SPCS that might constitute two distinct maps, rather than one (figure 8 in Kastner et al., 2007; figure 4 in Hagler et al., 2006). In the current study, we used eccentricity maps derived from pRFs to identify a foveal representation in SPCS. This, in turn, was critical to revealing the structure of SPCS1 and SPCS2. We believe that the pRF modeling technique we used here allowed us to more reliably measure topographic organization across all subjects.

Recent functional connectivity studies have observed two distinct clusters of connectivity patterns exactly at or near the junction of the PCS and superior frontal sulcus (Yeo et al., 2013; Yeo et al., 2011). Using functional connectivity, Wig et al. (2014) accurately identified the border between V1 and V2. More recently, researchers have shown that both SPCS and IPCS are divided into functional clusters that have a preference for visual or auditory space (Michalka et al., 2015). Future work should test if any of these clusters, either from resting-state connectivity or functional specialization, align to the border of SPCS1 and SPCS2 within individual subjects. This would likely provide great insight into the unique computations performed within the visual field map cluster in frontal cortex.

It is impossible to understand how a complex system processes information without first understanding how it represents information. Sensory systems are by far the most characterized systems in all of neuroscience, primarily due to our understanding of how they represent information, and our ability to use that knowledge to build models of the computations performed on those representations. This feat has proven far more difficult in higher-order cortical areas, as the further away a neuron gets from sensory receptors, the less that neuron's firing rate appears to correlate with quantifiable elements of the external environment. By identifying how higher-order areas, namely frontoparietal cortex, represent information, our current findings provide hope for understanding how higher-order brain regions process information and contribute to cognition and behavior.

## EXPERIMENTAL PROCEDURES

All of the data, along with the stimulus presentation and non-linear pRF model code is freely available on our website: http://clayspace.psych.nyu.edu.

### Subjects

Five neurologically healthy individuals (1 female, mean age 33, range 23-45) with normal or corrected-to-normal vision took part in the study. All subjects gave written informed consent before participating. All procedures were approved by the human subjects Institutional Review Board at New York University. Each subject completed one scanning session consisting of nine experimental runs.

### MRI Acquisition

MRI data were collected using a 3T head-only scanner (Allegra; Siemens) at the Center for Brain Imaging at New York University. Images were acquired using a custom four-channel phased array (NOVA Medical) placed over lateral frontal and parietal cortices, and a four-channel phased array placed beneath occipital cortex. Volumes were acquired using a T2*-sensitive echo planar imaging (EPI) pulse sequence [repetition time (TR), 2000 ms; echo time 30 ms; flip angle, 75°; 31 slices; 2.5 mm x 2.5 mm x 2.5 mm voxels]. T1-weighted anatomical images were collected at the beginning of each scanning session using the same slice prescriptions as the functional data. These were used to align the functional volume to a high resolution, whole brain anatomical scan. High-resolution T1-weighted scans (1 mm x 1 mm x 1 mm voxels) were collected for registration, segmentation, and display.

### Topographic mapping procedures

Observers performed a difficult discrimination task that required covertly attending to stimuli within bars of different widths that swept across the visual field in different directions. The total visual field was confined to a square, 24 deg on a side, with fixation in the center of the square. The length of the bar aperture was 24 degrees, and the width subtended 1, 2, or 3 degrees of visual angle. One of the three widths was used in any given 5-minute scan (similar to Winawer et al, 2010). Each bar aperture was split into three equal rectangular patches along its length. For example, a bar that swept from right to left was split into a top patch, center patch, and bottom patch. A bar that swept from top to bottom was split into a left patch, center patch, and right patch.

The bar aperture swept slowly but discretely across the visual field, from one end to the other. The steps were synchronized to the MRI acquisition (one step every 2 seconds), and the step size was 1.6 degrees. Each bar position defined one 2-s trial. For each trial, we asked subjects to select which of the two flanking patches of moving dots matched the direction of motion in the center patch. The dot motion in the center patch was 100% coherence so that its direction was unambiguous. The motion was along the length of the bar (up or down for vertical bars sweeping horizontally, and left or right for horizontal bars sweeping vertically). The direction of motion in one of the two flanking patches was matched to the center patch, and in the other flanking patch was opposite. In order to keep the discrimination task difficult, we used a two up one down staircase on the coherence value for the moving dots in the flanker patches. 124, 248, 372 Depending on bar size, each patch contained either 124 (1 degree bar), 248 (2 degree bar), or 372 (3 degree bar) dots (each 1/10 degree in size) moving at 1.6 degrees per second. The dot positions updated 60 times per second. For the flanker patches, the set of coherent dots was randomly re-selected on each frame update, so that no single dot moved continuously in one direction throughout a trial. Dots that were not coherent disappeared and were redrawn in a random location within the aperture in the subsequent frame. Stimuli were generated in MATLAB with the MGL toolbox and displayed on a screen in the bore of the magnet. Subjects viewed the display via a mirror mounted on the RF coil. Behavioral responses were recorded using a button box.

### MRI Preprocessing

T1-weighted anatomical scans were automatically segmented using Freesurfer (Dale et al., 1999). All fMRI analysis was performed using open source Matlab tool. The first three volumes of each functional run were removed to allow magnetization to reach a steady-state. Subsequent volumes were slice-time and motion corrected using tools by Kendrick Kay (https://github.com/kendrickkay/preprocessfmri). Data was then aligned to each individual subject’s T1-weighted anatomical image using a combination of vistasoft tools (http://github.vistalab/vistasoft) and Kendrick Kay’s align toolbox (https://github.com/kendrickkay/alignvolumedata). All subsequent fMRI analysis, including pRF analysis, was done using vistasoft. Functional scans for each individual experimental bar size (1, 2, and 3 degrees) were averaged together separately. Cortical surfaces were reconstructed at the gray/white matter border, and functional data (EPI time series) were projected to the gray matter voxels in the whole brain anatomy using trilinear interpolation. Data visualization projected model parameters from the gray voxels to a smoothed 3D mesh or flattened cortical representation.

### PRF analysis

We modeled response amplitudes for each voxel using a modified version of the pRF model described by Dumoulin & Wandell (2008) that incorporates a static power-law nonlinearity to account for nonlinear compressive spatial summation (CSS model, Kay et al., 2012). This model allows us to estimate an individual voxel’s receptive field center and size. Typically, the pRF model consists of an isotropic gaussian with four parameters: position (*x,y*), size (**σ**), and amplitude (**β**). The CSS model we employed adds an additional parameter, an exponent (*n*). This model is expressed formally as:

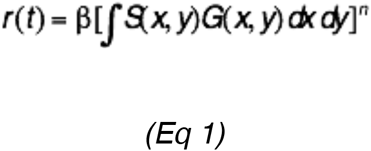

where *r(t)* is the voxel’s predicted response, *S* is the binary stimulus image, and *G* is an isotropic Gaussian expressed as:

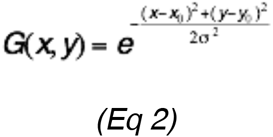

The original pRF fitting procedure described in Dumoulin and Wandell (2008) involved a two-stage fitting process: an initial coarse grid-fit followed by an exhaustive search fit using non-linear search optimization algorithms. Here, we used only the first stage (the grid fit), as it is more robust to noise and our goal was to map frontoparietal cortex where signals are much noisier and pRFs are larger than visual cortex. We fit model predictions to temporally decimated (2x) and spatially blurred (Gaussian kernel of 5mm width at half height) time series. Our gridded parameters included the same set of possible values in the solution described in Dumoulin and Wandell (2008), with the addition of the power law exponent (0.25, 0.5, 0.75, or 1). We interpolated solutions for voxels not included in the spatially blurred grid fit. We excluded voxels from further analysis in which the pRF model explained less than 10% of the variance of the time series, or which had pRF centers outside the limits of our visual display (12 degrees of visual angle) from further analysis.

### Model comparison

In order to compare nonlinear, linear, and contralateral model solutions, we cross-validated each by systematically solving the model on two-thirds of the data, and testing on the remaining one-third. As we used three different bar sizes in the experiment, each training data set contained six runs, with each bar size represented twice. The test data set included one run of each bar size. This was done three times so that all iterations of data shuffling under the constraint of having each bar size equally represented in a training or test sample were satisfied. Accuracy was defined as the variance explained of the model, and averaged for each model across the three different cross-validation iterations.

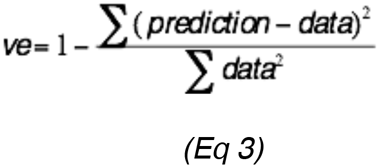

### Laterality index

We calculated a laterality index for each voxel based on *x0* and pRF size (*sigma*) parameters of the pRF model, similar to previous work (Sheremata and Silver, 2015). The only difference is that we define pRF size as 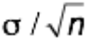 rather than just **σ** due to the compressive spatial summation (Kay et al., 2013).

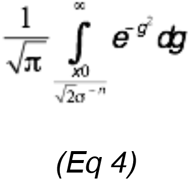

We also subtracted the lateralization index from 1 for left hemisphere voxels in order to compare between left and right hemispheres. This led to lateralization index values that were between 0 (completely ipsilateral) and 1 (completely contralateral).

## AUTHOR CONTRIBUTIONS

W.E.M., J.W., and C.E.C. designed research; W.E.M. performed research; W.E.M. analyzed data; W.E.M., J.W., and C.E.C. wrote the paper.

## ACKNOWLEDGEMENTS

This work was supported by National Institutes of Health Grant R01 EY016407 to C.E.C., R00 EY022116 to J.W., and the National Science Foundation Graduate Research Fellowship Program to W.E.M. We thank Lila Davachi, Martin Paré, Tommy Sprague, and David Heeger for comments on early versions of the manuscript.

